# Microbial habitability of early Mars lacustrine environments sustained by iron redox cycling

**DOI:** 10.1101/2021.09.07.459191

**Authors:** Rachel A. Moore, Christopher E. Carr

## Abstract

Several studies have reported new data on the estimated compositions of chemical components at Gale crater; however, there is still a lack of information regarding potential past support of biomass and detectable biomarkers of ancient life. In this study we evaluate microbial habitability of early Mars constrained by the recently reconstructed water chemistry at Gale. The modeled community is based on Fe-metabolizing bacteria with the ability to utilize solid-phase iron oxides (e.g., magnetite) as an electron source or sink. Our results illustrate the plausibility of a sustained community in Gale Lake and provides suggestions for future modelled and laboratory-based studies to further evaluate the past habitability of Mars, biosignatures and their preservation potential, and hidden metabolic potential.

**One Sentence Summary:** This work provides an existence proof of habitability on early Mars and demonstrates modeling processes by which the habitability of extraterrestrial environments can be explored quantitatively.

## 1. Introduction

The modern surface of Mars is analogous to Earth’s stratosphere: dry, cold, and inundated with DNA-damaging radiation. However, Mars possesses all the critical elements required for life including the essential elements (i.e., C, H, N, O, P, and S), and plausible sources of energy. It is possible that life could be present today in the Mars subsurface, beneath the cryosphere (Jones, Lineweaver and Clarke, 2011). Ancient Mars, with its higher abundance of liquid water, was arguably more hospitable to life as we know it, even while surface fluvial activity was likely transient, and temperatures generally cold (Wordsworth et al., 2021). For instance, Gale crater, a late Noachian to early Hesperian-aged crater (3.5-3.8 Gya), once harbored a liquid water lake (Grotzinger et al., 2015; Figure 1).

**Figure 1.**
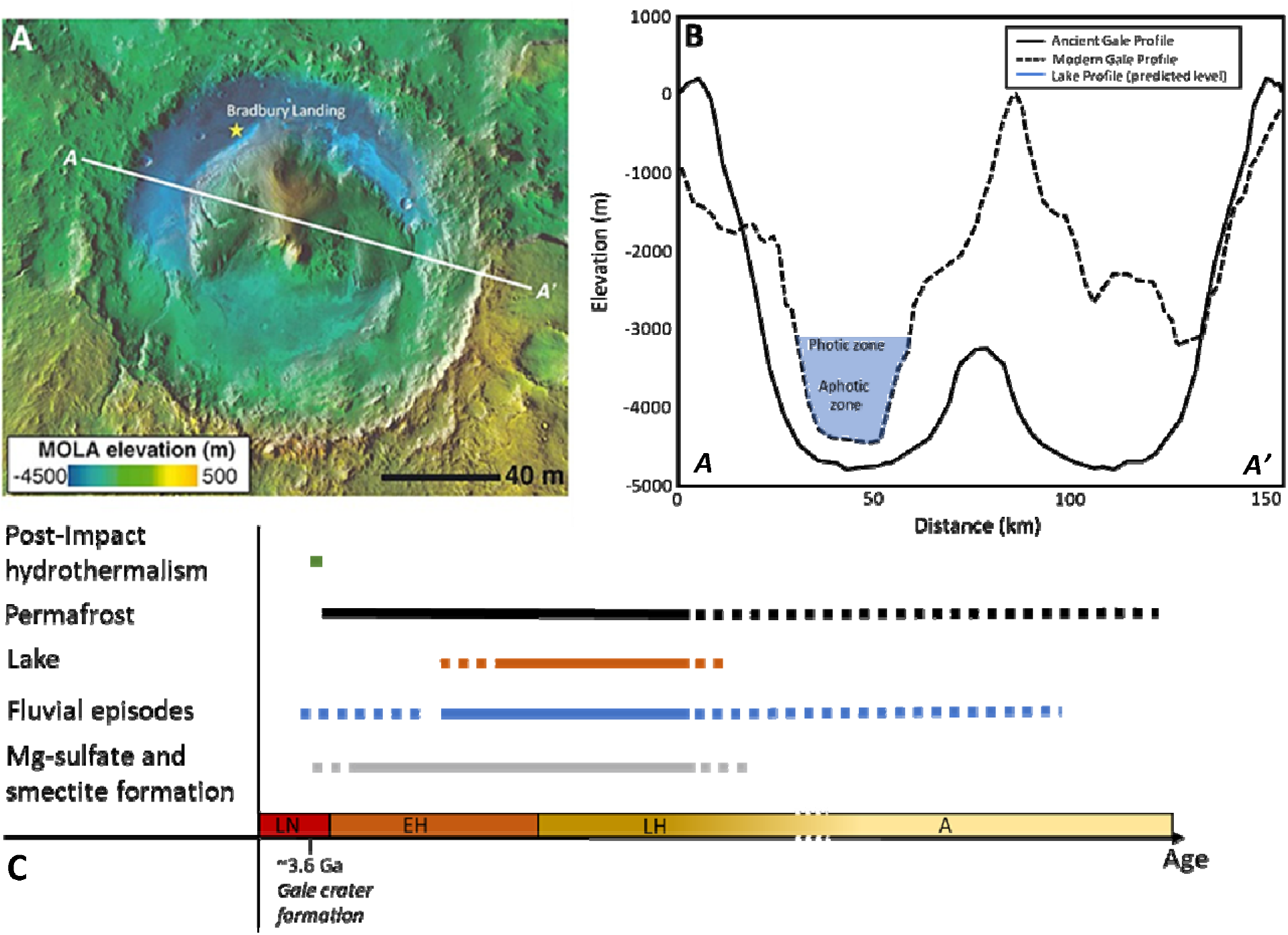
**A.** Aerial view of Gale Crater (137.4°E, −4.6°N) with star representing *Curiosity*’s Bradbury landing site. **B.** Profiles of Gale Crater: ancient (solid line) and modern (dashed line). The water level is not to scale. Modified from Borlina, Ehlmann and Kite, 2015. **C.** Geologic timeline of Gale Crater starting with its formation (LN: Late Noachian, EH: Early Hesperian, LH: Late Hesperian, A: Amazonian). Modified from Deit *et al.*, 2013.

Recently, samples collected by the Mars Science Laboratory rover *Curiosity* were used to estimate Gale Lake water chemistry (Fukushi et al., 2019). Major ions (Na^+^, K^+^, Mg^2+^, Ca^2+^, Cl^−^, SO_4_^2−^, HCO^3−^) and trace (e.g., Zn, Mg, Mn, Cu, Fe) components were constrained, in addition to pH and dissolved inorganic carbon (CO_2_). Taken together, this information provides rigorous constraints for modeling microbial habitability of Gale Lake.

Compatible microbial redox metabolisms have been suggested, however, they have yet to be experimentally assessed in an ancient Martian context (Cockell 2020). For example, several chemo-autotrophic and -heterotrophic redox couples would have been available for metabolism on the surface of Mars. Iron and sulfate ions have both been detected (Fukushi et al., 2019) and can be biologically reduced in conjunction with hydrogen. In addition, there is enough light energy and inorganic carbon to support anoxic photoautotrophy (Cockell, 2020). Phototrophy could have been productive in the "photic zone" of Gale Lake where solar energy was abundant and damaging radiation was attenuated (Cockell 2020; Figure 1 B).

This study approaches the possibility of maintaining a microbial community in Gale Lake established upon energy from light and iron redox cycling. The potential habitability of Mars and growing body of data assembled from *Curiosity* is the main reason for the choice of Gale crater as our study site. Our work proposes the use of *Geobacter sulfurreducens* and *Rhodopseudomonas palustris* as a simple biological community model to better understand habitability and production of detectable biomarkers in an inferred ancient lacustrine environments of Mars. Our work does not preclude other metabolisms, but provides an existence proof of habitability on early Mars and a process by which habitability of extraterrestrial environments can be explored quantitatively.

## 2. Materials and Methods

### 2.1. Development of genome-scale metabolic models

Previously reconstructed and validated genome-scale metabolic models of *R. palustris* (*iRpa940*; Alsiyabi, Immethun and Saha, 2019) and *G. sulfurreducens* (*iRM588*; Mahadevan *et al.*, 2006) were edited before their inclusion in community models. Several reactions for phototrophic growth in *R. palustris* were de-simplified based on the PNSB2011 model from Hädicke, Grammel and Klamt (2011) in order to more accurately track reaction fluxes. In addition, rxn01257, which produces precursors for folate and aromatic amino acids, was added to iRpa940 after gapfilling in KBase to allow for growth in Gale Lake minimal medium (Section 2.3). The enzyme associated with this reaction (i.e., aminodeoxychorismate synthase) has been identified in other Rhodopseudomonas spp. (e.g., #A0A1H8TD11).

The gapfilling process itself did not result in any additional added reactions to the model of *R. palustris*, however, several reaction directions were changed to be unidirectional (Supplemental file 1). Three reactions were added to *G. sulfurreducens* (i.e., rxn05291, rxn00197, rxn00257) during gapfilling. Reactions rxn00197, rxn00257, and rxn05291 are part of the Krebs cycle which is known to be present in *G. sulfurreducens* (Zhang et al., 2020). In both models, reactions were expanded upon to model iron redox cycling (Supplemental file 1), which has been previously demonstrated in this specific set of two organisms (Byrne *et al.*, 2015). Community models were created using COBRApy’s merge function with objective set to sum (Supplemental file 1). Separate cytoplasmic compartments were maintained by annotation of the reactions and metabolites for each organism. The community model is represented in Figure 2.

**Figure 2.**
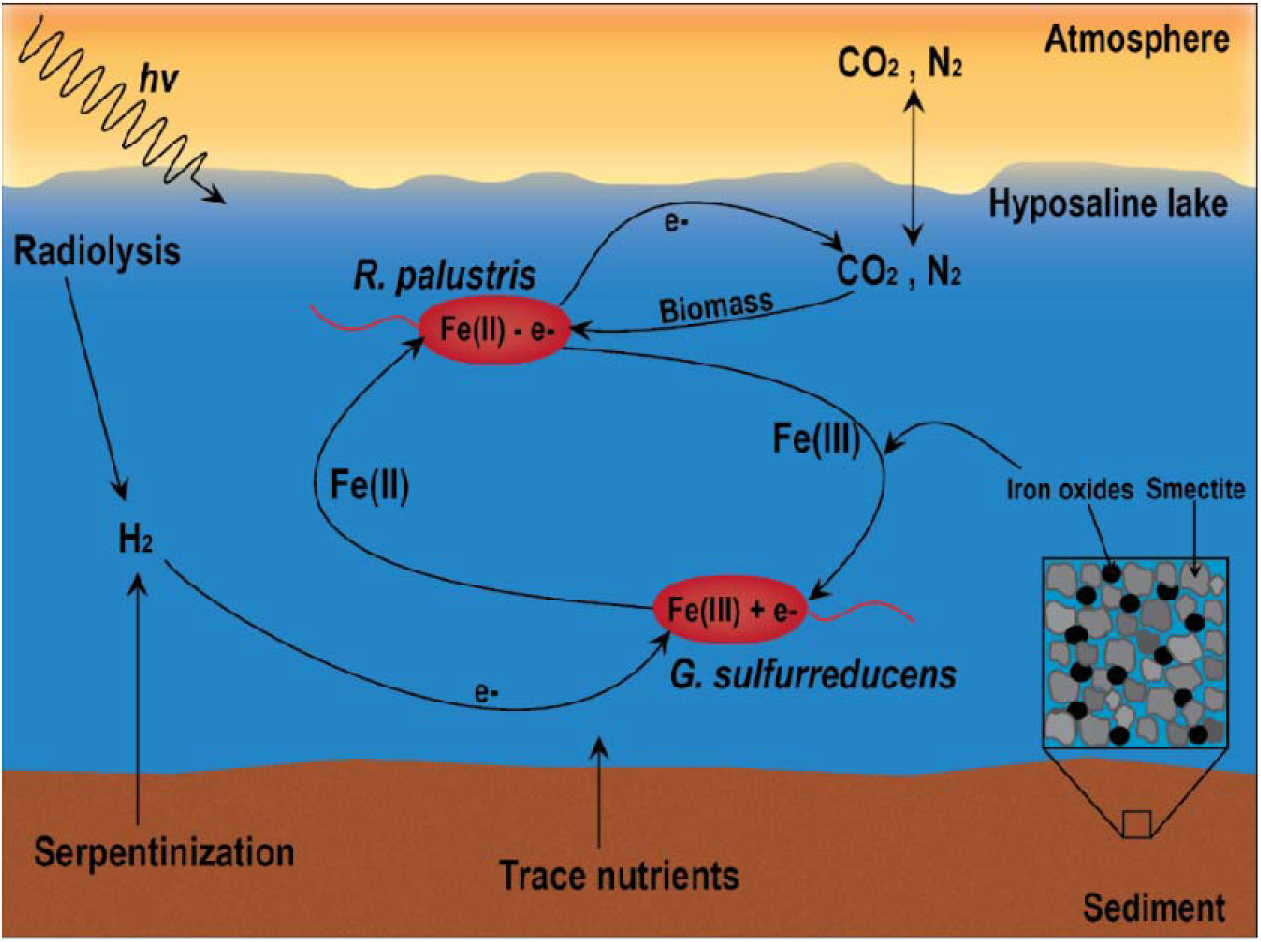
Model of Gale Lake emphasizing iron redox cycling phenomena.

### 2.2 Construction of the diurnal community model

A diurnal model was constructed, based on the diuFBA model from Knies et al. 2019, using the community model of *R. palustris* and *G. sulfurreducens* in section 2.1 as a scaffold. Briefly, the individual metabolic models were duplicated, and a separate extracellular space was defined (“night”, e2). Photon flux was only allowed in the “day” model (e0). An ATP maintenance reaction (rxn00062) was added to both organisms so that they would have to realistically expend energy even when not prioritizing growth. The reaction rates were based upon how much energy each organism spends on non-growth-associated maintenance (Klamt, Schuster and Gilles, 2002; Mahadevan *et al.*, 2006).

The day and night models were connected via transfer reactions that allow the “storage” and use of any metabolite, apart from photons and protons, in a different period other than when it was produced (Equation 1).

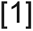

The transfer reactions are bidirectional, as a microorganism can theoretically store metabolites from night and use them during the day, as well as store from day to use at night. This is illustrated in Figure 3.

**Figure 3.**
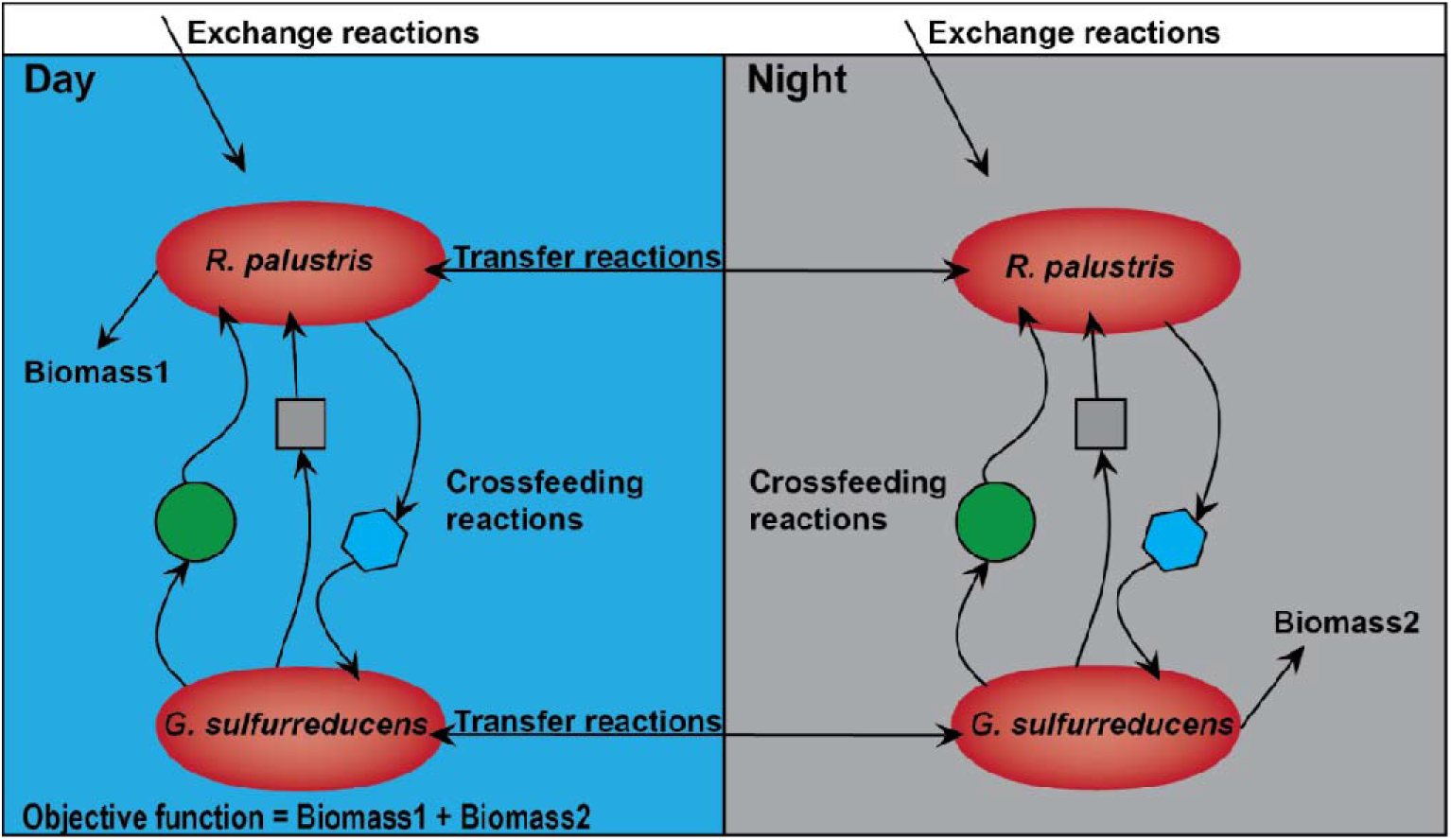
Schematic of the diurnal community model.

A single biomass reaction occurs for *R. palustris* in “Day”, and in “Night” for *G. sulfurreducens* to account for organism growth based upon when it would be prioritized (Byrne *et al.*, 2015). Both reactions are summed to serve as the overall objective function (Fig. 3). Flux values were integrated over 12 hours for both light and dark periods to estimate the total amount of each metabolite during one diurnal cycle (24-hour period) as in equation 2:

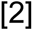

where v is the flux of a given metabolite (mmol gDW^−1^ h^−1^), and Δt is the duration of either the light or dark period (12 h). The result, Δc, corresponds to the molar amount of a given metabolite produced in the light or dark period (mmol gDW^−1^; Knies *et al.*, 2015).

### 2.3 Formulation of a Gale Lake minimal medium

An *in-silico* medium (Table 1) was simulated based on the methods of Marinos *et al.* (2020) and the composition of reconstructed water chemistry and evidence of trace minerals (Fukushi *et al.*, 2019; Payré *et al.*, 2019). Copper and zinc deposits within Gale crater were likely adsorbed onto Mn-oxides and thus insoluble (Payré *et al.*, 2019). For this reason, the bounds for copper and zinc were based on the optimal rate of import from the external environment for the community model. The remaining metabolites (Table 1) were treated as soluble within the lake system. The proton flux (H+) reflects the neutral pH of Gale Lake proposed by Fukushi *et al.* (2019). Concentrations of cobalt and magnesium were calculated from the *in situ Spirit* and *Opportunity* rovers’ measurements reported in Payré *et al.* (2019) as follows:

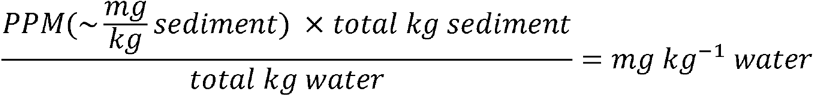

**Table 1.** Estimated chemical composition of Gale Lake and corresponding bounds utilized in the modelled growth medium.

| Metabolite / SEED compound ID | Concentration (mol L<sup>-1</sup>) | Flux upper bound (mmol gDW<sup>-1</sup> h<sup>-1</sup>) |
| --- | --- | --- |
| Ca<sup>2+</sup> / cpd29674 | $2.4\text{-}4.5 \times 10^{-2} *$ | 24 - 45 |
| Cl<sup>-</sup> / cpd00099 | $1.1\text{-}2.5 \times 10^{-1} *$ | 110 - 250 |
| Co<sup>2+</sup> / cpd00149 | $1.0\text{-}5.9 \times 10^{-6} ****$ | $1.0\text{-}5.9 \times 10^{-3}$ |
| CO<sub>2</sub> / cpd00011 | $2.3\text{-}41 \times 10^{-3} *$ | 2.3 - 41 |
| Cu<sup>2+</sup> / cpd00058 | $1.0 \times 10^{-7} ***$ | $1.0 \times 10^{-4}$ |
| Fe<sup>2+</sup> / cpd10515 | $1.2\text{-}58 \times 10^{-4} *$ | $1.2\text{-}58 \times 10^{-1}$ |
| H<sup>+</sup> / cpd00067 | $1.0 \times 10^{-6} **$ | $1.0 \times 10^{-3}$ |
| H<sub>2</sub> / cpd11640 | $1.2 \times 10^{-2} ***$ | 12 |
| H<sub>2</sub>O / cpd00001 | 1.0 | 1000 |
| K<sup>+</sup> / cpd00205 | $1.4\text{-}4.4 \times 10^{-3} *$ | 1.4 – 4.4 |
| Mg / cpd00254 | $3.5\text{-}6.0 \times 10^{-2} *$ | 35 - 60 |
| Mn<sup>2+</sup> / cpd00030 | $5.3\text{-}100 \times 10^{-7} ****$ | $5.3\text{-}100 \times 10^{-4}$ |
| N<sub>2</sub> / cpd00528 | $130\text{-}220 \times 10^{-5} *****$ | $1300\text{-}2200 \times 10^{-2}$ |
| Na<sup>+</sup> / cpd00971 | $9.4\text{-}12 \times 10^{-2} *$ | 94-120 |
| PO<sub>4</sub><sup>3-</sup> / cpd00009 | $5.2\text{-}18000 \times 10^{-7} ***$ | $5.2\text{-}18000 \times 10^{-4}$ |
| SO<sub>4</sub><sup>2-</sup> / cpd00048 | $4.4\text{-}7.2 \times 10^{-2} *$ | 44-72 |
| Zn<sup>2+</sup> / cpd00034 | $1.0 \times 10^{-7} ***$ | $1.0 \times 10^{-4}$ |

\* Data from Fukushi *et al.* (2019) total dissolved components of calcium, chloride, carbon dioxide, iron, hydrogen, potassium, magnesium, sodium, and sulfate, respectively.
\*\* Calculation based on data from Fukushi *et al.* (2019)
\*\*\* Concentration and flux based on model requirements.
\*\*\*\* Calculation based on data from Lanza *et al.* (2014) and Berger *et al.* (2019)
\*\*\*\*\* Calculation based on Von Paris *et al.*, (2013) and Navarro-Gonzalez *et al.* (2018)

The total mass (6.67×10^6^ kg) of sediment in Gale was derived from the estimated lake area (2×10^10^; Fukushi *et al.*, 2019) and the average bulk density of Martian sediment (3×10^3^ kg m^−3^). Total water mass (3×10^15^ kg) was the lower bound estimated in Fukushi *et al.* (2019). The upper limit concentration of N_2_ was calculated using the ideal gas law, a maximum partial pressure of 0.3 bar (Von Paris *et al.,* 2013; Navarro-Gonzalez et al 2018), a temperature of 273.15 K (Fukushi *et al.*, 2019) and a Martian atmospheric mass of 2.5×10^16^ kg (*Mars Fact Sheet*, 2021). The lower bound for N_2_ was based on the optimal rate of import from the external environment. Water was allowed to freely enter the extracellular environment at 1000 mmol gDW^−1^ h^−1^.

For models with photoautotrophic growth, photons were allowed to enter the external environment at a rate of 100 mmol gDW^−1^ hr^−1^ (9.7 μE m^−2^ s^−1^).

### 2.5 Flux balance analysis

The metabolic flux distribution of the individual, community, and diurnal models was calculated using flux balance analysis or dynamic flux balance analysis. The specific growth rate was used as the objective function (function to be maximized) in all simulations except for those where maximum yields of hopanoids were estimated as in Orth, Thiele and Palsson (2010). All simulations were performed using the Python programming language (version 3.8) in conjunction with open-source GLPK software (https://www.gnu.org/software/glpk/), and COBRApy (Ebrahim *et al.*, 2013), pandas (Reback *et al.*, 2020), and NumPy (Harris *et al.*, 2020) libraries.

## 3. Results

### 3.1 Genome-scale metabolic models

Table 2 shows the specific growth rate for each model on the different types of substrates with or without light. Although *R. palustris* is capable of chemotrophic growth in complete media (Alsiyabi, Immethun and Saha, 2019), it fails to grow in the modelled medium without the presence of light (Table 2) or a source of organic carbon. Further, all other models grow slowly at low levels of metabolite flux in the medium (0.001 h^−1^; Table 2). At higher levels of metabolite flux, community growth is roughly two orders of magnitude faster than all other models (Table 2). For example, growth of the community in the presence of light (1.031 h^−1^) increased ~2000% as compared to growth of the individual models combined (i.e., 0.005 and 0.032 h^−1^ or 0.037 h^−1^ total).

**Table 2.** Specific growth rates of the genome-based models under the specified conditions and constrained to either the low or high bounds of medium metabolite flux.

| Electron donor/acceptors | Input carbon source | Light? | Model | Specific growth Rate ($\text{h}^{-1}$) |
| --- | --- | --- | --- | --- |
| $\text{H}_2 / \text{Fe}$ | $\text{CO}_2$ | NA | *G. sulfurreducens* | 0.001–0.005 |
| $\text{Fe} / \text{CO}_2$ | $\text{CO}_2$ | - | *R. palustris* | 0.00–0.00 |
| $\text{Fe} / \text{CO}_2$ | $\text{CO}_2$ | + | *R. palustris* | 0.001–0.032 |
| $\text{H}_2, \text{Fe} / \text{Fe}, \text{CO}_2$ | $\text{CO}_2$ | + | Community | 0.001–1.031 |
| $\text{H}_2, \text{Fe} / \text{Fe}, \text{CO}_2$ | $\text{CO}_2$ | - | Community | 0.001–1.012 |

NA: Not applicable

Syntrophic interactions were investigated and are presented in Table 3. Both *R. palustris* and *G. sulfurreducens* produce metabolic products consumed by the other member. For example, carbon sources (acetate and citrate) were produced and exported to the extracellular environment where “cross-feeding” can occur. Acetate produced by *G. sulfurreducens* was consumed by *R. palustris* at a rate of 142.5 mmol gDW^−1^ h^−1^ when grown in the higher range of medium fluxes. Likewise, citrate was produced at a rate of 63.29 mmol gDW^−1^ h^−1^ by *R. palustris* and consumed by *G. sulfurreducens* (Table 3). Iron (Fe^2+^ and Fe^3+^) was consumed and produced at the same rate (4.38 mmol gDW^−1^ h^−1^) between the microbes and the extracellular environment (Table 3).

**Table 3.** Modelled interactions between community members^*^.

| Metabolite | *R. palustris* (reaction ID) | *G. sulfurreducens* (reaction ID) |
| --- | --- | --- |
| Acetate | -0.0053 / -142.5 (rxn05488) | 0.0053 / 142.5 (rxn05488) |
| Fe<sup>2+</sup> | -0.022 / -4.38 (rxn37614_2) | 0.022 / 4.38 (FERCYT) |
| Fe<sup>3+</sup> | 0.022 / 4.38 (rxn37614_2) | 0.022 / -4.38 (FERCYT) |
| Citrate | -0.0037 / 63.29 (rxn05211) | 0.0037 / -63.29 (rxn05211) |
| CO<sub>2</sub> | 0.0072 / -94.55 (rxn05467) | -0.22 / 55.41 (rxn05467) |

<sup>\*</sup> Fluxes are shown in mmol gDW<sup>-1</sup> h<sup>-1</sup> for low and high range of medium inputs with light. Negative values indicate metabolite consumption, whereas positive values indicate production.

#### 3.1.1 Chemolithoautotrophic metabolism of G. sulfurreducens

Flux balance analysis revealed an unexpected hidden autotrophy, namely that *G. sulfurreducens* was capable of carbon dioxide fixation through the reverse Krebs cycle (Fig. 4). Here citrate (Table 3) was used (rxn00257) to start the cycle. CO_2_ was fixed at two steps, from succinyl-CoA to 2-oxoglutarate (Low-high = 0.27-43.31 mmol gDW^−1^ h^−1^; rxn00197) and from 2-oxoglutarate to isocitrate (Low-high = 0.075-30.3 mmol gDW^−1^ h^−1^; rxn00198).

**Figure 4.**
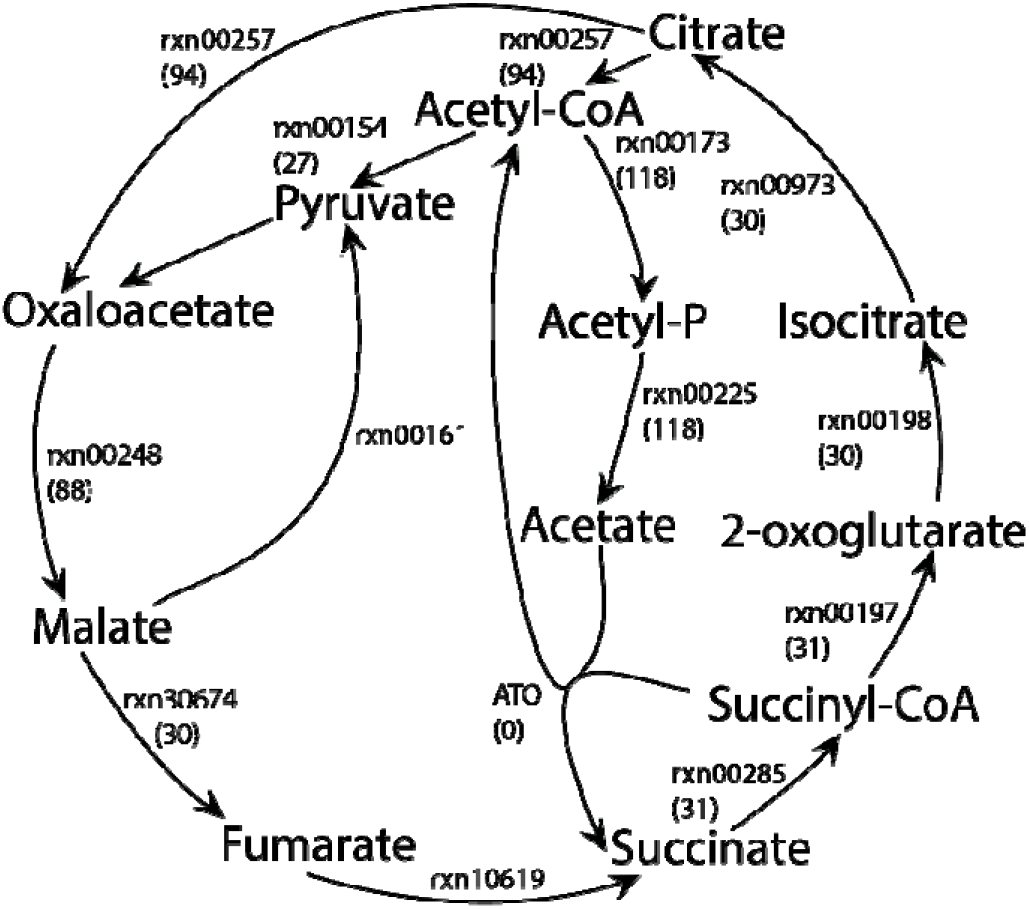
The reverse Krebs cycle of *G. sulfurreducens* uncovered by flux balance analysis. Numbers in parentheses refer to reaction fluxes (mmol gDW^−1^ h^−1^, rounded to nearest whole number). The arrows refer to the direction in which each reaction is taking place.

### 3.2 Dynamic flux balance analysis

Dynamic growth of the community model based on nitrogen (Fig. 5), the limiting metabolite, was used to calculate the total biomass supported per mL of Gale Lake in the photic zone. Nitrogen was considered limiting for growth in this context as all other necessary metabolites were capable of being recycled or produced following initial input. Independent of initial media metabolite flux (i.e., high or low), 0.23 grams of biomass were produced per gram of N_2_ (Fig. 5). Depending on the amount of nitrogen present (1.3 – 2.2 mmol N_2_ L^−1^), Gale Lake could have maintained 10^3^-10^7^ cells per mL of brine, assuming cells are 10^−12^ grams in weight on average (Bratbak and Dundas, 1984; see supplemental materials).

**Figure 5.**
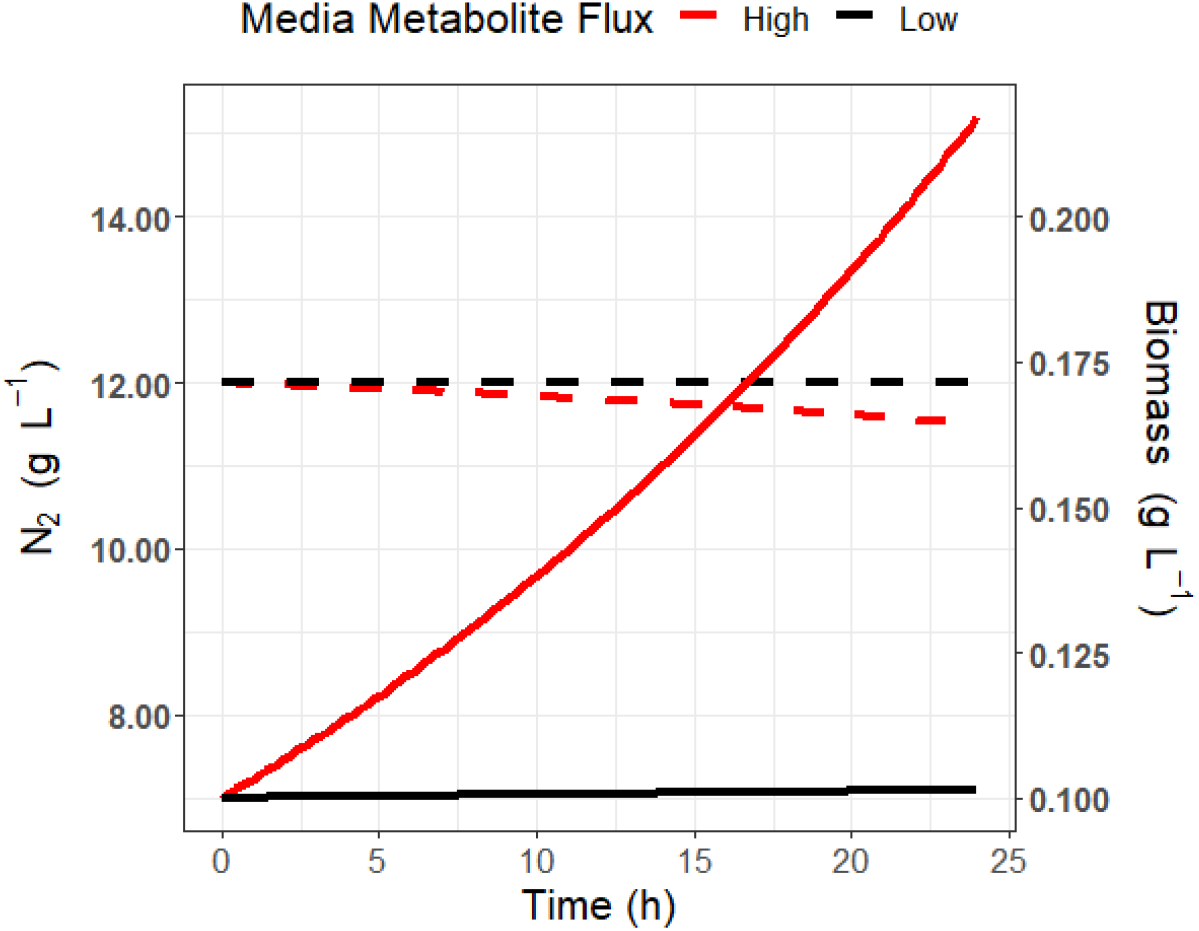
Production of community biomass (solid lines) and consumption of N_2_ (dashed lines) with low (black) or high (red) flux of metabolites in the medium.

### 3.3 Hopanoid production

To quantify hopanoid production, the production of precursor farnesyl diphosphate was used as the objective function. Nitrogen was limited to 1 mmol gDW^−1^ h^−1^ to predict hopanoid maximum yields per mmol of limiting substrate (N_2_). Farnesyl diphosphate is produced at 0.001 mmol per mmol of N_2_. If, as estimated, Gale Lake contained 13 mmol N_2_ L^−1^, we predict ~0.01 mmol of hopanoid was produced per liter in the photic zone. This translates to a range of 2×10^−1^ - 2×10^−2^ grams of hopanoid kg^−1^ of Gale Crater sediment assuming a photic zone depth of 100 m and a sediment thickness of 1-10 m (see supplemental materials), assuming a preservation event (such as settling after a mass death event) of 100% of the cells and uniform mixing of settled material within the sediment.

### 3.4 Diurnal community model

The overall growth rate of the diurnal community model was twofold greater than the traditional community model (with light, 2.062 h^−1^ vs 1.031 h^−1^). In addition, *G. sulfurreducens* produced more biomass (1.998 h^−1^) as compared to *R. palustris* (0.0645 h^−1^). Overall, concentrations of metabolites (i.e., consumed and secreted) did not change significantly between periods of light and dark except for nucleic acids (7.61×10^3^ vs. 1.27×10^4^) and vitamins and cofactors (9.74×10^3^ vs. 1.26×10^4^) for *R. palustris* (Fig. 6, Supplemental File 2). Transfer reaction fluxes indicated that both organisms prioritized the period in which they were growing. For instance, 5 out of the 8 metabolite categories were associated with a negative transfer flux for *R. palustris* indicating transfer from night to day (Fig. 7). Likewise, 6 out of the 8 categories were associated with positive flux values for *G. sulfurreducens* (Fig. 7).

**Figure 6.**
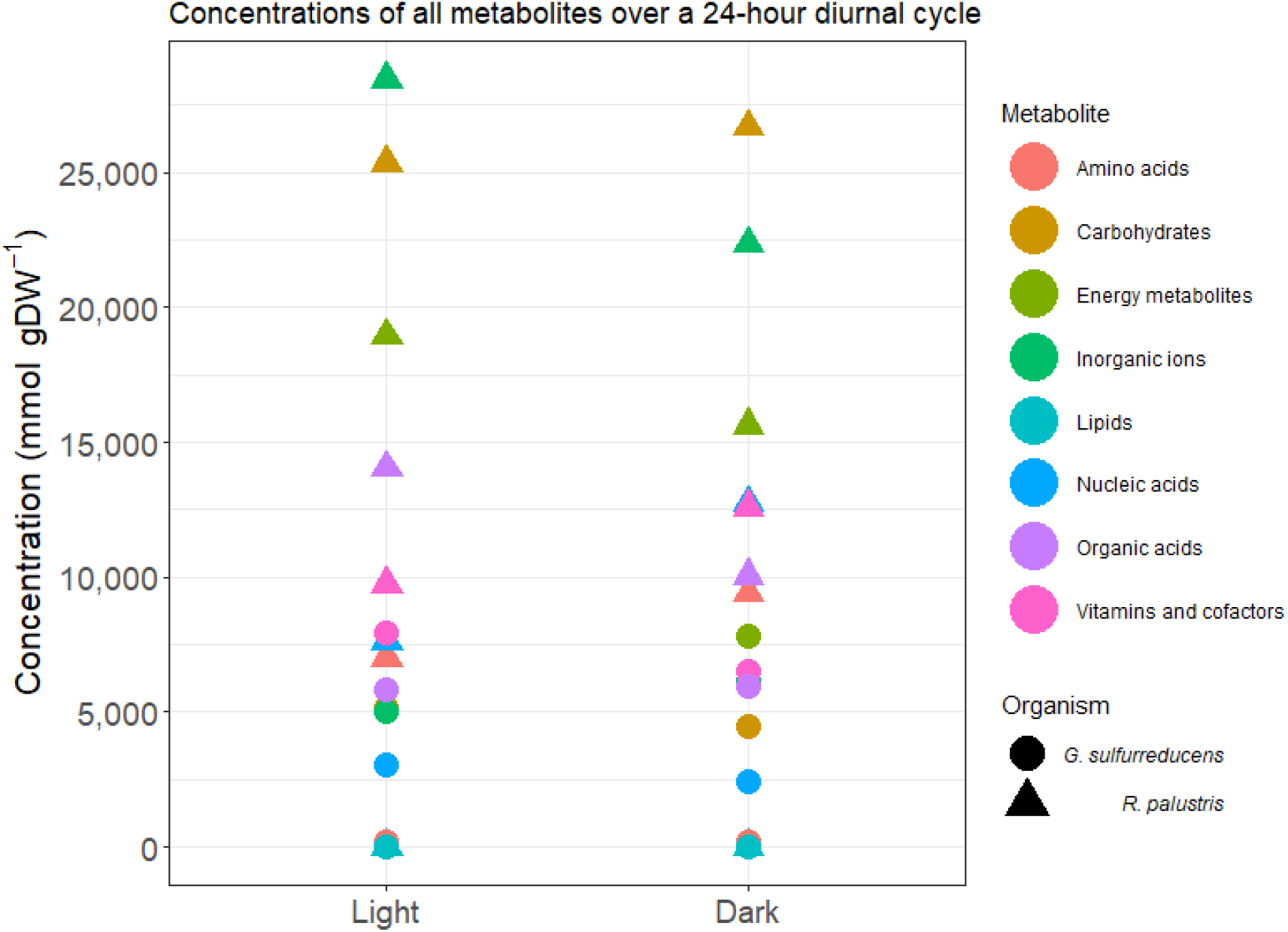
Metabolite concentrations summed by category for 12-hour periods of light and dark. Circles and triangles represent *G. sulfurreducens* and *R. palustris*, respectively.

**Figure 7.**
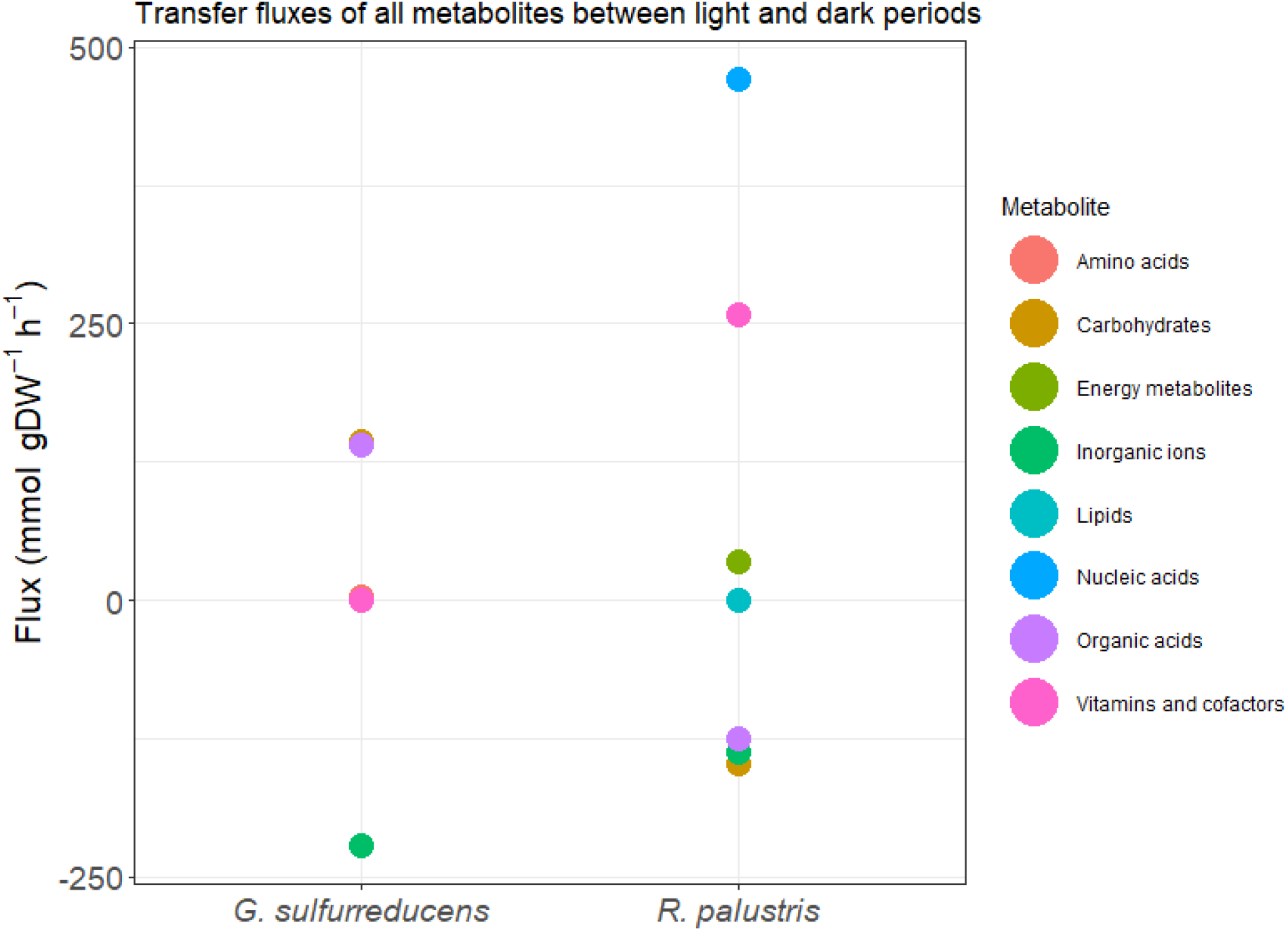
Metabolites transfer fluxes between light and dark summed by category. Negative values indicate transfer from night to day, and positive values indicate transfer from day to night.

#### 3.4.1 *R. palustris* transfer reaction fluxes and ATP consumption

The nucleic acid and energy metabolite flux profiles for *R. palustris* (Fig. 7) were governed mainly by thymidine triphosphate (TTP) and guanosine triphosphate (GTP), respectively. TTP and GTP were transferred from light to dark where they served as phosphate donors to produce ATP from ADP (Fig. 7). Carbohydrates (i.e., 1,3-Bisphospho-D-glycerate and glycerone) were primarily transferred from night to day in *R. palustris* (Fig. 7). These were ultimately used to generate NADP and ATP through the production of phosphate. Inorganic ions like thiosulfate were transferred to daytime where they were used to create acetate and homocysteine, and to reduce cytochrome c3. Out of the organic acids, oxaloacetate was transferred between night and day (Fig. 7). Oxaloacetate was primarily used to power the daytime Krebs cycle.

#### 3.4.2 G. sulfurreducens transfer reaction fluxes and ATP consumption

Carbohydrates, inorganic ions, and organic acids were transferred at nonzero fluxes between night and day for *G. sulfurreducens* (Fig. 7). Of the carbohydrates, acetyl phosphate and citrate are transferred to night where they are used to make ATP. Specifically, citrate is used to power the reverse Krebs cycle (Fig. 4). The inorganic ion flux is dominated primarily by phospshate. Phosphate is transferred from night to day, where it is used to produce acetylphosphate. Acetylphosphate is used to generate ATP in both day and night. Any acetate (organic acids) produced is secreted to the extracellular space.

## 4. Discussion

In this work we evaluate the microbial habitability of early Mars’ lacustrine environment constrained by reconstructed Gale Crater water chemistry and sustained by the redox cycling of iron. In this context, we used *G. sulfurreducens* and *R. palustris* as model organisms because they have been shown to effectively and anoxically use solid-phase iron oxides (Byrne *et al.*, 2015). Although these specific organisms have been successfully co-cultured, this is not a known natural association. However, similar microbial communities have been isolated *in natura*. For example, an Fe(II)-oxidizing photoferrotroph, *Candidatus Chlorobium masyuteum*, and a putative Fe(III)-reducing bacterium, *Candidatus Pseudopelobacter ferreus*, were recently co-enriched from the ferruginous Brownie Lake in Minnesota, USA (e.g., Lambrecht *et al.*, 2021). Acetate and formate detected in the chemocline of Brownie Lake suggests that the redox cycling of Fe occurs between these organisms, but it has not yet been confirmed. Until recently, it was thought that no single organism was capable of both Fe(II)-oxidation and Fe(III)-reduction. It is now understood that the bacterium *Rhodoferax* strain MIZ03 can utilize iron as an electron donor and acceptor (Kato and Ohkuma, 2021).

In an evolutionary context, Alpha- and Delta-proteobacteria like *R. palustris* and *G. sulfurreducens*, respectively, probably did not arise on Earth until well after (<2-3 GYA; Marin *et al.*, 2017; Wang and Luo, 2021) Gale crater was formed in the Late Noachian (Fig. 1C). If we assume an origin of life took place ~4 GYA (i.e., either on Earth or Mars; Cantine and Fournier, 2017; Carr, 2021) and that the hypothesis of common ancestry between Earth and any possible Mars life is true (Velasco, 2018), these specific metabolisms would likely have emerged after Gale hosted a lake (~3.8 to 3.1 GYA; Hurowitz *et al.*, 2017). The modeled organisms are reasonable for testing early Mars habitability if we assume an earlier second genesis occurred. If instead we assume life evolved independently (i.e., if the hypothesis of common ancestry is false) on Earth, these metabolisms also could have emerged in time to inhabit Gale Lake.

It is important to note that a potential caveat of constraint-based modeling is that it does not account for temperature. Growth rates would be affected by changes in temperature but not growth yield. In addition, our models assumed a constant light intensity. Daily light fluctuations have been accounted for in previous models (e.g., Sarkar *et al.*, 2019), however, this granular approach becomes exponentially complex for multi-species models and was not necessary for our assessment. Further, as *R. palustris* is a chemoautotroph and primary producer, we did not supply the models with additional carbon sources other than CO_2_. Organic carbon is present on Mars (Freissinet *et al.*, 2015; Eigenbrode *et al.*, 2018) but its distribution is not known.

In the present study, the individual models were capable of slow growth (Table 2) on the modelled medium but were predicted to grow faster together as a community, indicating syntrophy (Table 3). For instance, iron was effectively cycled by being reduced and oxidized at the same rate (Table 3). Like the findings of Byrne et al. (2015), these results indicate that iron does not become biologically unavailable to either organism unless perhaps something were to physically limit access (e.g., biofilm formation). We also found that the citrate secreted by *R. palustris* (Table 3) was consumed and used by *G. sulfurreducens* in the reverse Krebs cycle (Fig. 4), illustrating genome-scale metabolic modelling as a discovery tool. It has recently been confirmed that *G. sulfurreducens* can utilize this cycle *in vitro* to fix CO_2_ (Zhang *et al.*, 2020). Such hidden autotrophy could also have implications for planetary protection if microbial metabolism is used to guide planetary protection requirements.

A diurnal community model was constructed to further assess cross-feeding interactions and metabolite storage during periods of light and dark (Fig. 3). Previous models have been used to illustrate the circadian growth of other microbes (e.g., cyanobacteria), but they have not been modelled in a community setting (e.g., Sarkar *et al.*, 2019). We found that both organisms transferred metabolites to periods in which they prioritized growth (Fig. 7). For example, *R. palustris* prioritized carbohydrate use during the day to release phosphate, which is necessary to produce ATP. Additional ATP was produced at night via nucleoside-diphosphate kinases to feed the non-growth maintenance reaction. Conversely, *G. sulfurreducens* primarily transferred carbohydrates like citrate and acetyl phosphate to produce ATP for growth at night.

In the context of early Mars, life, if it existed there, would have likely been present in the form of a community. Microorganisms in nature are rarely found in isolation, with only one known example to date (i.e., *Candidatus Desulforudis audaxviator*, Chivian *et al.*, 2008). This is because communities are better able to cope with changes in temperature, humidity, and nutrients (Palková, 2004). The syntrophic interactions identified here highlight the benefits of a community, which could allow life to adapt to varying environmental conditions just as shifts occur in our simple diurnal model. Beneficial community interactions might also have enabled life to achieve higher biomass and biosignature production than might otherwise have been possible.

Our calculations show that Gale Lake could have maintained a concentration of cells (~10^3^-10^7^ cells mL^−1^) similar to modern lakes and oceans. For example, modern terrestrial ocean waters and saltern brines have been shown to support 10^4^-10^6^ cells per mL (Bar-On, Phillips and Milo, 2018); 10^4^-10^7^ Klempay *et al.*, 2021) depending on depth and other variables. Terrestrial ocean and lake sediments have also been shown to harbor 10^4^-10^9^ cells per cm^3^ (Kallmeyer *et al.*, 2012; Wurzbacher *et al.*, 2017).

In addition to predicting the habitability and biomass density of early Mars, it is of particular interest to calculate the potential densities of biomarkers (i.e., molecular fossils) that could be used to detect ancient life. For example, certain membrane lipids like bacteriohopanepolyols or hopanoids can persist for billions of years in the terrestrial rock record (Brocks *et al.*, 2005). The hopanoids synthesized by many bacteria are largely resistant to degradation because of their pentacyclic carbon skeleton, and can be detected through several analytical techniques (e.g., high-performance liquid chromatography, liquid/gas chromatography–mass spectrometry; Belin *et al.*, 2018). Here, both *R. palustris* (Welander *et al.*, 2009) and *G. sulfurreducens* (Härtner, Straub and Kannenberg, 2005) are known to produce biomarker hopanoids under anaerobic conditions in culture. Our model predicts that hopanoids could have been produced at a concentration (~10^−1^-10^−2^ g kg^−1^ Gale sediment; see supplemental materials) well above the limit of detection for miniaturized GCMS systems (~1 PPM, Coy et al., 2011) in the case where all cells in the photic zone at some steady state timepoint settled and were evenly distributed into Gale Lake sediments and were 100% preserved. While this represents an upper bound for a single such event, such events could have occurred repeatedly, due to dust storms or impact event-driven dust events blocking the sunlight required for photosynthesis. Future work could link steady state hopanoid (or other biosignature) production to deposition, preservation, and diagenesis, to yield refined estimates in support of *in situ* life detection.

## 5. Conclusions

Here we have demonstrated that Gale Lake could have provided the energy necessary to sustain a simple and syntrophic microbial community. It must be noted that our models and calculations contain several assumptions. For instance, Gale Lake would not have had an even distribution of light, or of any ‘metabolite’ for that matter, throughout its depths. It is also pertinent to note that the densities of all lake components would have changed greatly over time with evaporation (e.g., Rapin *et al.*, 2019). This would also inevitably lead to reduction in hopanoid concentration potentially rendering it more difficult to detect. Nevertheless, these data provide guidance for Mars habitability studies. Future research should further develop and confirm candidate biosignatures *in vitro* and using other scenarios of habitability on early Mars and on other worlds.

